# Cerebellar tDCS alters the perception of optic flow

**DOI:** 10.1101/2021.02.15.431326

**Authors:** Jean-François Nankoo, Christopher R Madan, Omar Medina, Tyler Makepeace, Christopher L. Striemer

**Author notes:** Corresponding author: Jean-François Nankoo, PhD, Krembil Research Institute, Toronto Western Hospital Toronto, Ontario, Canada, M5T 2S8.

## Abstract

Studies have shown that the cerebellar vermis is involved in the perception of motion. However, it is unclear how the cerebellum influences motion perception. tDCS is a non-invasive brain stimulation technique that can reduce (through cathodal stimulation) or increase neuronal excitability (through anodal stimulation). To explore the nature of the cerebellar involvement on large-field global motion perception (i.e., optic flow-like motion), we applied tDCS on the cerebellar midline while participants performed an optic flow motion discrimination task. Our results show that anodal tDCS improves discrimination threshold for optic flow perception, but only for left-right motion in contrast to up-down motion discrimination. This result was evident within the first 10 minutes of stimulation and was also found post-stimulation. Cathodal stimulation did not have any significant effects on performance in any direction. The results show that discrimination of planar optic flow can be improved with tDCS of the cerebellar midline and provide further support for the role of the human midline cerebellum in the perception of optic flow.

## Introduction

Until recently, the role of the cerebellum was thought to be primarily confined to motor control and the coordination of movement. However, we now know that the cerebellum is also involved in a variety of non-motor functions such as memory, emotion, and perception [1,2]. Of particular interest to the current work, mounting evidence suggests that the integrity of the cerebellum is important for motion perception (see [1] for review). Anatomical, lesion, and non-invasive brain stimulation (NIBS) studies suggest an important role of the cerebellar vermis in motion perception. However, there remains many questions regarding the role of the vermis on motion perception. For example, it is unclear whether the vermis is involved in large-field global motion perception, such as optic flow, and whether the mechanisms by which the vermis influences motion perception are inhibitory or excitatory.

Early evidence provided by Nawrot and Rizzo [3,4] showed that patients with acute midline cerebellar lesions are significantly impaired when performing a motion discrimination task. Specifically, the tasks involved required participants to identify the global direction (up, down, left, right) of random dot stimuli (RDS) while the signal to noise ratio was varied to obtain threshold. The RDS were displayed within a small 4° × 4° square region. Patients with midline cerebellar lesions, but not those with lateral lesions, showed elevated thresholds. No differences were found between patients with hemispheric cerebellar lesions and healthy controls (see also [5]). The authors suggest that the performance of patients with midline lesions is akin to patients with lesions in the medial temporal (MT) area, a cortical region critical for motion perception [6,7]. Building on the work of Nawrot and Rizzo [3,4], Cattaneo et al. [8] showed that transcranial magnetic stimulation (TMS), a NIBS technique, significantly impaired performance when applied over the cerebellar vermis during a motion discrimination task. Similarly to Nawrot and Rizzo [3,4], the participants in Cattaneo et al. [8] viewed RDS within a small display that subtended 4.3° × 4.3° of visual angle. The participants were required to indicate whether the global motion pattern moved to the left or the right. The results showed that disruption of the cerebellar vermis impaired performance, thus confirming the involvement of the cerebellar vermis on motion perception. However, Cattaneo et al. [8] used a ‘virtual lesion’ TMS protocol that does not provide any information about the mechanism (i.e., inhibitory or excitatory) by which the cerebellar vermis affects motion perception.

From an anatomical perspective, the cerebellum has multiple connections to areas that are highly specialized to process motion information. For instance, the cerebellum is connected to the cortical motion areas medial temporal (MT)/medial superior temporal area (MST) through the pontine nuclei as well as the thalamus [9–11]. Cells in MT/MST have been shown to respond to RDS and direct stimulation of MT neurons in rhesus monkeys has been found to bias the perception of motion direction [12,13]. Functional brain imaging and lesion studies have substantiated the importance of V5/MT+, the homologue of MT/MST, in motion perception in the human brain [7,14]. Antal et al. [15] showed that cathodal tDCS on V5 improved motion discrimination (using RDS) performance. They argued that cathodal stimulation acts to suppress the neuronal noise that ultimately allows the signal to be more easily detected. In addition to the connectivity with V5, the cerebellum also receives input from the accessory optic system, a subcortical visual pathway present in all vertebrates that is specialized for processing optic flow [16]. Optic flow is the motion pattern on the retina that arises as we move through the environment and provides the brain with information about body and eye movements which are critical for controlling balance and gait, and for determining direction of heading [17]. Although it is known that optic flow is processed in the flocculo-nodular lobe, it is currently unclear whether modulating excitability of the vermis can influence the perception of optic flow.

NIBS techniques are useful for establishing the function of brain structures as well as for inducing plasticity [18,19]. One such technique is transcranial direct stimulation (tDCS), a method that involves passing a small amount of current through the scalp. This current is thought to increase, via anodal (+) stimulation, or decrease, via cathodal (−) stimulation, activity of neurons near the stimulated area by changing the resting-membrane potential of neurons [20,21]. In this study, we investigated the role of the cerebellum in the perception of optic flow by applying tDCS to the midline cerebellum. If the cerebellar vermis is important for processing optic flow, then altering activity levels in the cerebellum using active (i.e., anodal (+) or cathodal (−)) tDCS should alter discrimination thresholds compared to a sham stimulation condition.

## Materials and Methods

### Experiment 1

Participants consisted of 16 right-handed undergraduate students from MacEwan University (Male = 3; Female = 13; mean age = 21.44; SD = 2.25). All the participants had normal or corrected-to-normal vision. The participants received course credit and/or monetary compensation of $10 per hour for participating in the study. All the participants were naïve to the purpose of the experiment. The experiment received ethics approval from the MacEwan University Research Ethics Board and was conducted in accord with the Code of Ethics of the World Medical Association (Declaration of Helsinki).

### Apparatus

The stimuli were generated on a Dell Optiplex 9020 with an Intel i7 processor using MATLAB (The MathWorks Inc., Natick, MA) and the Psychophysics Toolbox [22,23]. The stimuli were displayed on a white wall using Epson PowerLite Home Cinema 2030 projector with a refresh rate of 60Hz. Participants were seated comfortably in a dark room at a viewing distance of approximately 40 cm to the display. Responses were collected using a keyboard.

### Stimuli and Design

The stimuli consisted of RDS with varying levels of coherence. The dots were white on a black background and subtended a visual angle of 82° × 53°. The dot density in the display was 1%. Each dot subtended a visual angle of 0.05° × 0.05°, and moved at a velocity of 9.3°/s per frame (3 pixels at each refresh) and lasted five frames (i.e., limited lifetime [24,25]) before being repositioned in a random location within the display. In the left/right (LR) trials, all the signal dots moved either leftward or rightward. In the up/down (UD) trials, all the signal dots moved either upward or downward. Regardless of signal direction, each noise dot moved to a randomly selected direction and maintained that trajectory for the duration of the trial. Each stimulus was presented for a total duration of 600 ms (60 Hz image update rate), and the inter trial interval was set at 500ms (Fig. 1).

**Fig. 1.**
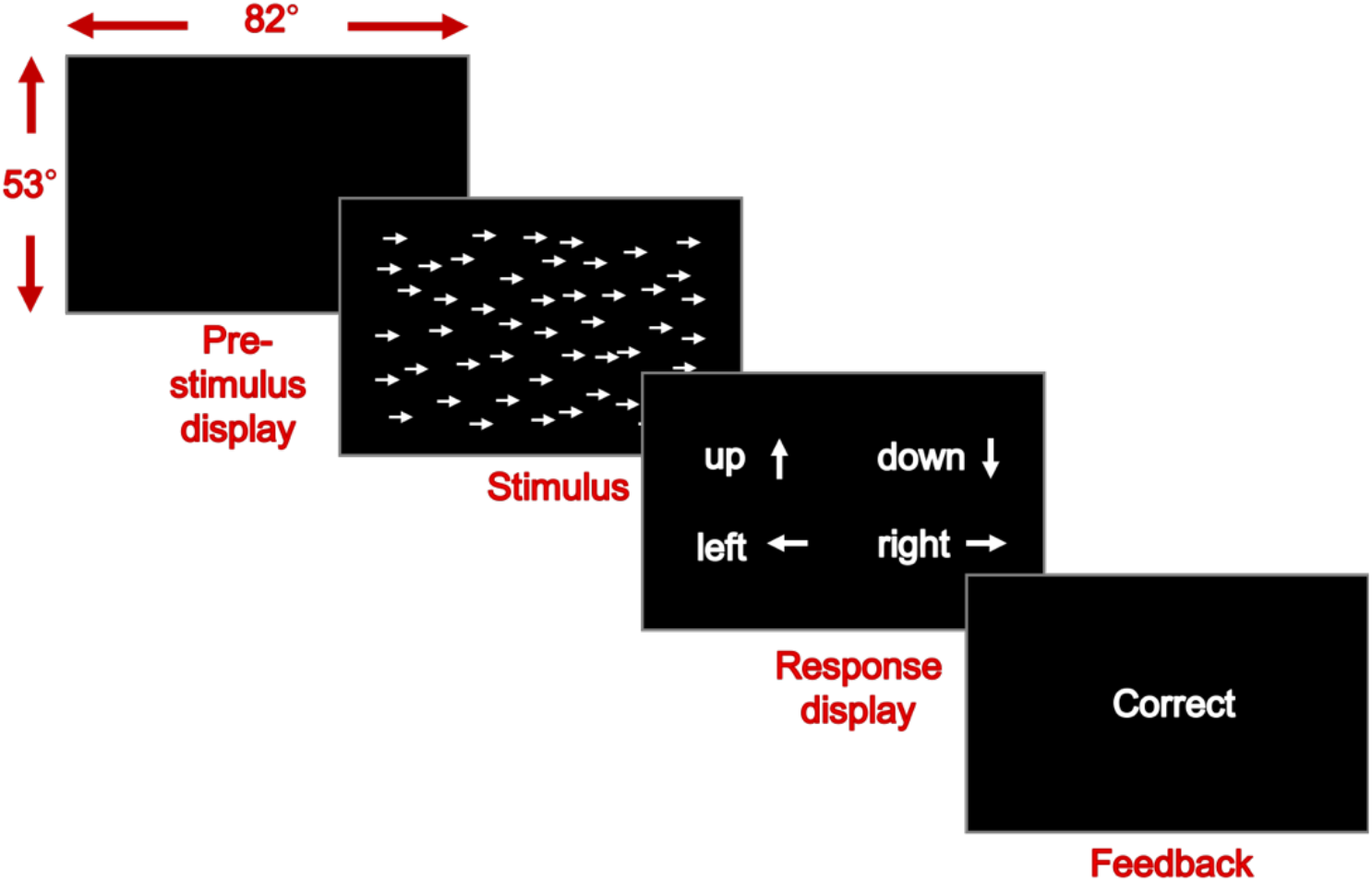
Illustration of the stimulus presentation protocol. First, a blank display (Pre-stimulus display) is presented and after 500 ms, a random dot stimuli is presented for 600 ms. Immediately following the stimulus display, the participant is prompted to choose the key that correspond to the direction of motion of the random dot arrow. A visual feedback is given (i.e., “Correct” or “Incorrect”) following the response. The visual screen sustained a visual angle of 82° × 53°.

### Transcranial Direct Current Stimulation

For the tDCS procedure, participants were fitted with saline-soaked sponge tDCS electrodes. The active electrode (anode (+) or cathode (−)) was placed on the scalp along the midline of the cerebellum by measuring 2 cm below the inion. During active tDCS, a 2mA current was administered for 20 minutes with a 5cm × 5cm saline soaked sponge electrode [18,26] via a battery powered Mind Alive Oasis Pro tDCS stimulator (Mind Alive, Edmonton, AB; http://mindalive.com/). This generated a current density of .08mAa/cm^2^. The current was ramped up over 10s at the beginning of stimulation and ramped down over 10s at the end of stimulation. The reference electrode was placed on either the right or left shoulder (randomized across participants) with a 10 cm × 10 cm saline-soaked sponge electrode. These stimulation parameters are well within the established safety guidelines for tDCS administration in humans [27,28]. Using these stimulation parameters, we constructed a current distribution map across the cerebellum using the SimNIBS software package [29]. The model revealed that the current distribution was largely centered on the cerebellum with little spread to V1 or V5/MT (Fig. 2).

**Fig. 2.**
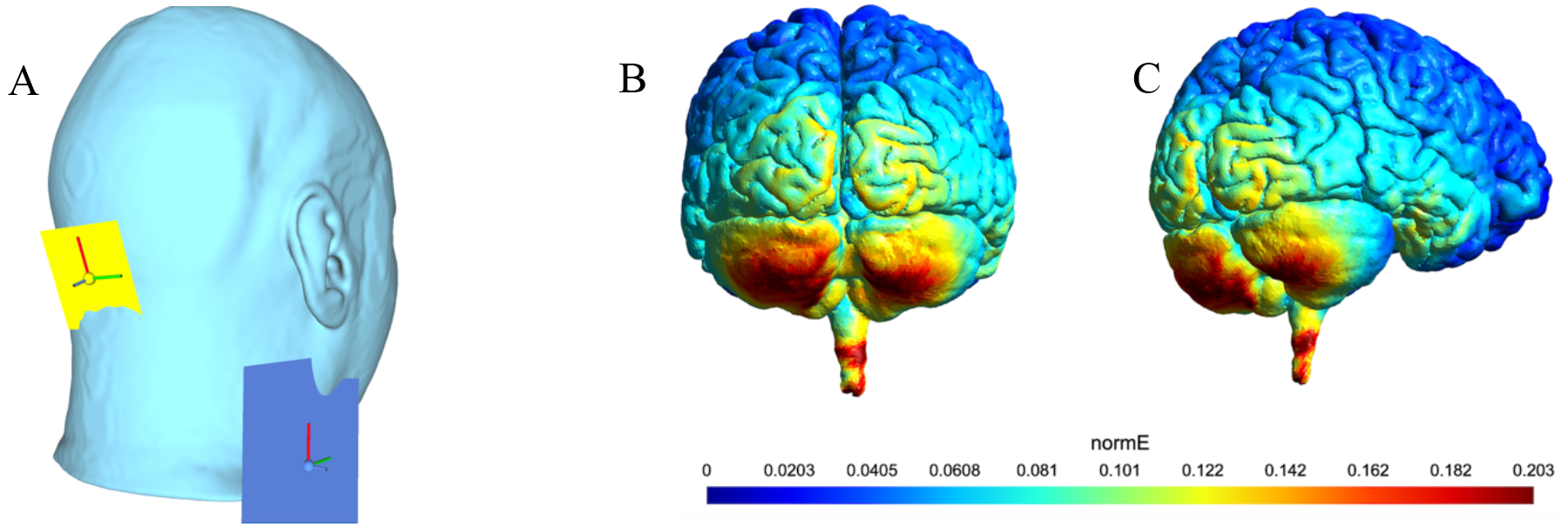
Transcranial direction current stimulation (tDCS) simulation software, SimNIBS v3.0.5[29] was used to create a map of the current across the cerebellum, mapped to a template brain model. Panel A shows the electrode montage. The active electrode was modeled as a 5×5cm simple whole-surface electrode positioned at midline, positioned at the scalp 2 cm below the inion. The reference electrode was modeled as a 10×10cm simple whole-surface electrode mounted on the junction of the lower neck and shoulder. Simulated stimulation with 2.0 mA of current at the active electrode yielded an estimated electric field (NormE) across the ipsilateral cerebellar hemisphere as seen in panel B (posterior view) and panel C (posterolateral view).

The experiment was run as a within-subject design, such that each participant completed each of the three stimulation conditions (anodal (+), cathodal (−), and sham) on a different day with all conditions completed within one week. The order of the stimulation conditions was randomized across participants.

During the sham stimulation condition (tested on a separate day), participants underwent the exact same procedure as the anodal and cathodal condition with the exception that the stimulator remained on (so it appeared to be working), but the current was turned off (unbeknownst to the participant) after 30s. This ensured that participants felt the same initial mild itching and tingling sensations that were experienced during the anodal and cathodal stimulation conditions.

### Procedure

For each session, participants were given a practice run of ten trials (5 UD and 5 LR). The participants were then presented with stimuli of varying coherence and were not told whether it was a LR or UD trial. The participants responded by pressing one of the four arrow keys on a keyboard (↑↓←→). This means that on each trial, regardless of the motion direction, the participant had a 25% chance of randomly picking the correct response key.

A Thresholding block followed the practice session in which the thresholds for UD and LR motion for each participant were obtained using the QUEST estimates based on 80 trials for each direction [30]. The Thresholding block was followed by a Stimulation block. The Stimulation block consisted of trials in which the stimuli were shown at threshold levels and the participants were stimulated with anodal (+), cathodal (−), or sham tDCS. The percentage correct for UD and LR were recorded. The first and second ten minutes of the stimulation were considered as two blocks (Stimulation 1 and 2) to allow for an analysis of stimulation duration. Afterwards, the Post-Stimulation block consisted of the same setup as the Stimulation block with the exception that the tDCS sponge was removed from the scalp and no stimulation was given. The trials for both motion directions (i.e., LR and UD) were randomized across trials for a total of 160 trials (80 trials for each motion direction) for each block. Participants completed a total of 480 trials per session. Feedback was given on the screen by displaying the word “Correct” or “Incorrect” as shown in Fig. 1.

### Data Analysis

The detection thresholds were determined by a maximum likelihood procedure using the QUEST adaptive staircase procedure [30]. In the QUEST procedure, the participant’s psychometric function is assumed to follow a Weibull distribution [31] and coherence levels are based on responses in previous trials. Threshold for both motion directions were obtained within the same block. The trials were randomized, but each threshold measurement was independent. Threshold was taken as the proportion of dots to achieve 72.5% accuracy as we found this number to provide accurate estimates from QUEST. All statistical analyses were conducted using JASP (JASP Team, 2020; Version 0.14). Effects were considered significant based on an alpha level of .05, and multiple pairwise comparisons were corrected using the Holm-Bonferroni method [32].

## Results

The mean detection threshold obtained prior to each stimulation session is shown in Fig. 3. Mean coherence threshold for UD motion obtained on days of anodal stimulation was .11 (*SD* = .06). On cathodal days, the mean threshold also .10 (*SD* = .05). On sham days, the mean coherence threshold was also .10 (*SD* = .04). Mean coherence threshold for LR motion were as follows: On days of anodal stimulation the mean was .10 (*SD* = .05), on cathodal days the mean was .09 (*SD* = .04), and on sham days the mean was .08 (*SD* = .05). A two-way repeated-measures ANOVA of stimulation day (i.e., threshold obtained on anodal, cathodal and sham days) and motion direction (UD, LR) yielded a non-significant main effect of stimulation day (*F*(2, 30) = 1.38, *p* = 0.27, partial *η*^2^ = .08) and motion direction (*F*(1, 30) = 1.53, *p* =.24, partial *η*^2^ = .09). The interaction of stimulation day and motion direction was also non-significant (*F*(2, 30) = 0.24, *p* = .79, partial *η*^2^ = .02). Given that there were no significant differences in pre-stimulus thresholds across the different testing sessions, the accuracy for each testing condition was taken as the proportion of correct answers above or below threshold performance (72.5%).

**Fig. 3.**
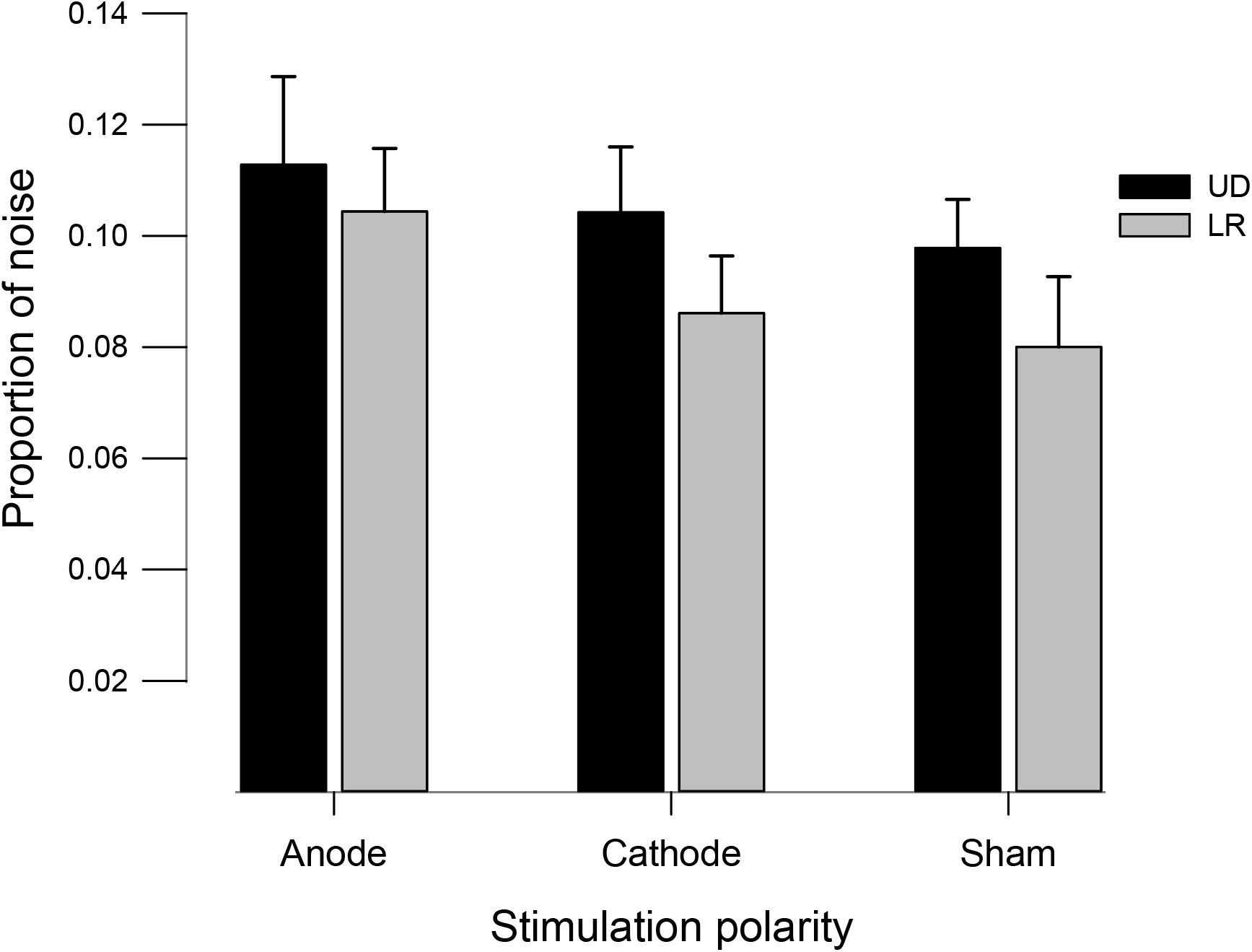
The mean detection threshold (72.5% accuracy) obtained prior to each stimulation session in Experiment 1. UD = Up-Down motion discrimination threshold; LR = Left-Right motion discrimination threshold. Error bars represent SEM.

The main analyses were performed on the mean accuracy differences expressed as a change from the threshold criterion (72.5% accuracy) for each stimulation duration (10 minutes stimulation, 20 minutes stimulation, and post-stimulation), stimulation polarity (sham, anodal, and cathodal), and motion direction (UD and LR). These data are summarized in Supplementary Table 1. A repeated-measures ANOVA resulted in a significant main effect of duration (*F*(2, 30) = 27.73, *p* < .001, partial *η*^2^ = .65), and stimulation polarity (*F*(2, 30) = 4.20, *p* = .025, partial *η*^2^ = .22). However, no main effect of motion direction (*F*(1, 15) = 1.27, *p* = .28, partial *η*^2^ = .08) was found. A significant interaction of stimulation polarity and motion direction was also detected (*F*(2, 30) = 3.53, *p* = .04, partial *η*^2^ = .19; Fig. 4). No other interactions were significant (all *p*s > .05).

**Fig. 4.**
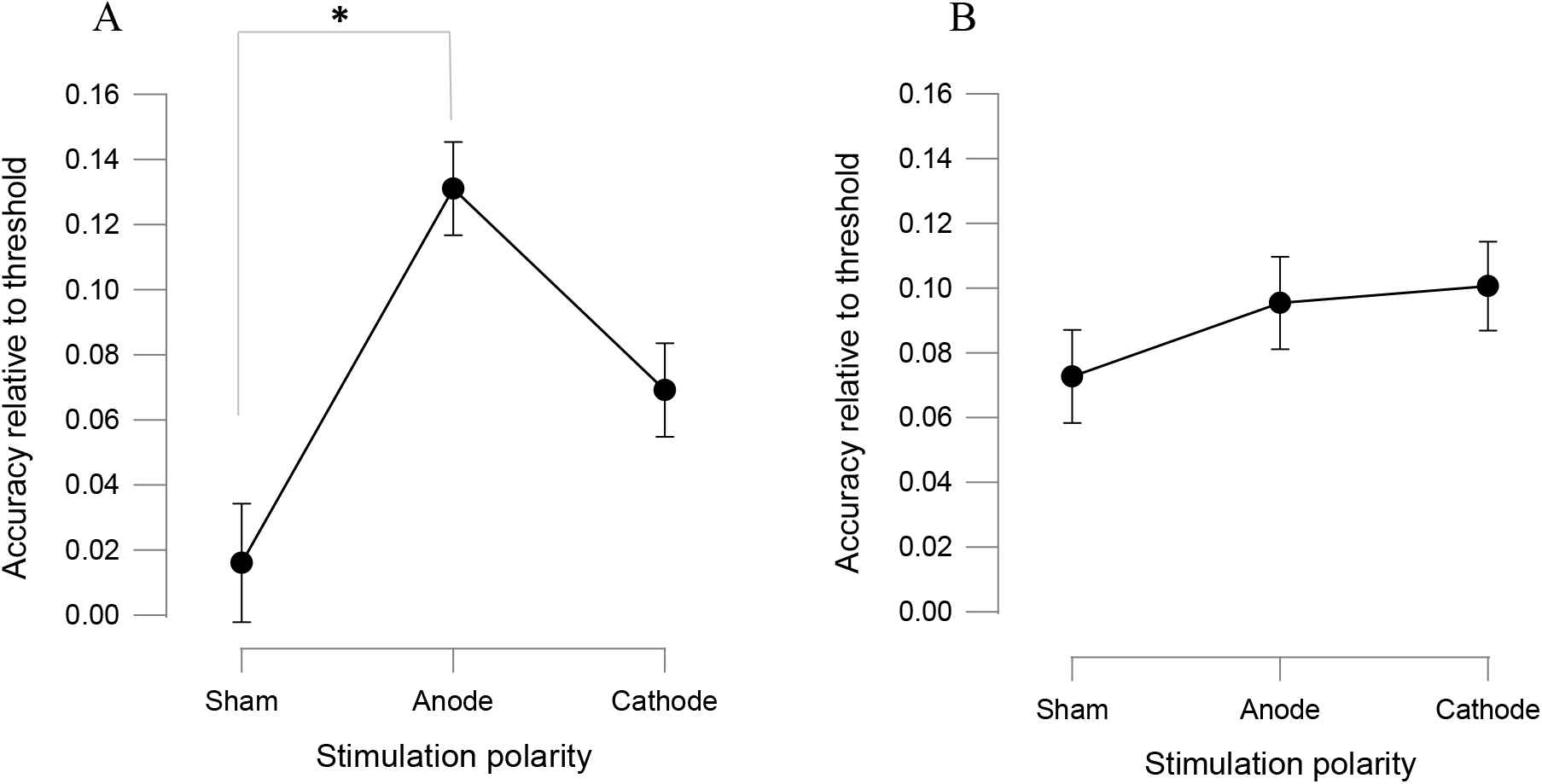
Interaction effect of stimulation polarity (Sham, Anode, and Cathode) and motion direction (UD and LR) on the mean accuracy difference from the threshold criterion (72.5% accuracy). **Panel A** shows the effects of stimulation polarity on LR motion, whereas **panel B** shows the effects of stimulation polarity on UD motion. Positive values represent an improvement in performance, negative values represent a decrease in performance, and zero values represent no change from baseline. Error bars represent SEM and **\*** represents *p* < .05.

To explore the interaction of stimulation polarity and motion direction, we ran separate RM ANOVAs for each motion direction. For UD motion, no significant effect of stimulation polarity was detected (*F*(2, 30) = 0.66, *p* = .56, partial *η*^2^ = .04; Figure 4B). However, a significant effect of stimulation polarity was found for LR motion (*F*(2, 30) = 5.93, *p* = .007, partial *η*^2^ = .28; Figure 4A). Holm corrected post hoc pairwise comparisons showed that LR performance was significantly better in the anodal condition compared to sham (*M*_anode_ = .13, *SD* = .09; *M*_sham_ = .016, *SD* = .12; (*t*(15) = 3.44; *p* = .005, *d* = .86), but no significant differences emerged between anodal and cathodal stimulation (*M*_cathode_ = .07, *SD* = .09; (*t*(15) = 1.85; *p* = .15, *d* = .46), and between cathode and sham conditions (*M*_sham_ = .02, *SD* = .11; (*t*(15) = 1.59; *p* = .15, *d* = .40).

Given previous reports of an effect of cerebellar TMS discrimination between up and down motion (e.g., Cattaneo et al., [8]), we carried an additional experiment to eliminate the possibility that the difference in tDCS effects we found on UD and LR were not simply due to the difficulty of our procedure. In the follow-up experiment we used the exact same tDCS procedures but with a two-alternative forced choice paradigm using only UD motion in order to rule out the possibility that presenting both LR and UD motion within the same session may have been too taxing for the participant and perhaps biased attention to the LR discrimination.

### Experiment 2

The method for Experiment 2 was identical to Experiment 1 with the exception that we only tested UD. Consequently, the participants were instructed to choose either the up or down arrow key, and there was a 50% chance of identifying the correct direction. The participants (*n* = 16; Male = 7; Female = 9; mean age = 21.94; SD = 4.17) in Experiment 2 were naïve and did not participate in Experiment 1.

The mean coherence threshold for UD motion obtained on days of anodal stimulation was .12 (*SD* = .11), whereas on days of cathodal stimulation the mean threshold was .08 (*SD* = .04). For sham stimulation, the mean threshold was .09 (*SD* = .09). A repeated-measures ANOVA on the change in accuracy relative to thresholds resulted in a significant main effect of duration (*F*(2, 30) = 5.33, *p* = .01, partial *η*^2^ = .26), but no effect of stimulation polarity (*F*(2, 30) = 0.67, *p* = .52, partial *η*^2^ = .04). The interaction between duration and stimulus polarity was found to be non-significant (*F*(4, 60) = 0.40, *p* = .81, partial *η*^2^ = .03).

## Discussion

We investigated the role of the midline cerebellum on large field motion perception (i.e., simulated optic flow) using tDCS, a NIBS technique that has been shown to modulate cortical excitability and influence behaviour (see for a review [33]). First, we established the proportion of noise needed for each participant to achieve a performance of 72.5% (i.e., threshold) on each testing day. Given that no significant differences were detected in the proportion of dots required to perform at threshold, we analyzed performance based on changes from 72.5% accuracy. Specifically, we found that performance on a motion discrimination task was improved, relative to threshold, during and immediately following anodal tDCS of the midline cerebellum. However, the improvement was only found when discriminating between left-right motion (Fig. 4A, while no effect was found when discriminating between upward and downward motion (Fig. 4B). In addition, no statistically significant effect of cathodal tDCS was found for either left-right or up-down motion discrimination. Although the difference between sham and cathodal tDCS did not achieve statistical significance, we did observe a slight improvement of performance in the left-right discrimination condition from cathodal stimulation (Fig. 4A). We also found that the effects of anodal stimulation on left-right motion discrimination performance was significantly improved throughout the experiment (i.e., at 0-10min, 10-20min, and post) as there was no interaction with time. In a second experiment we confirmed that midline tDCS has no effect on up-down optic flow discrimination. This result highlights the specificity of the effects of tDCS to left-right optic flow. Overall our results show that tDCS can improve optic flow perception post-stimulation which is indicative of plasticity. Because optic flow is intrinsically linked to gait, our results indicate that tDCS may be promising for improving the perception of optic flow in patients with gait deficits such as patients with Parkinson’s disease or stroke survivors [34,35].

However, because tDCS has a low spatial resolution, the results of our study should be interpreted with caution. Based on our modelling of the electrical fields, it is likely that we stimulated parts of the cerebellar hemispheres, in addition to the vermis (see Fig. 2 and [36]). The hemispheres, like the vermis, have been shown to be involved in motion processing [37, 38]. However, in contrast to the hemispheres, the vermal lobules have been linked specifically with processing optic flow and are related to input from the accessory optic system (AOS), and project directly, and indirectly, to the cerebellar vermis [39, 40, 41, 42]. Cells within AOS have been shown to have receptive fields about 40° or more ([43, 44, 45, 46]). Therefore, it likely that the larger stimuli (i.e., greater than 40°) would engage the optic flow system in comparison to the smaller stimuli. Research from Baumann and Mattingley [37, 38], who found cerebellar hemisphere involvement in motion perception, have used stimuli that were substantially smaller compared to the current study (Baumann & Mattingley [37] = 26° × 20°; current study= 82° × 53°). The size differences in stimuli between the current study and those used in Cattaneo et al. [8], may also explain the differences between the two studies (Cattaneo et al. [8] = 4.3° × 4.3°). Unlike Cattaneo et al. [8] we did not observe any effects on up-down motion discrimination. Thus, the results of Cattaneo et al., [8] and Baumann & Mattingley [37, 38], along with the data from the current study, provide converging evidence for the involvement of the cerebellum in the perception of motion, as has been shown in the cerebral cortex [47]. However, Cattaneo [8] and Baumann and Mattingley’s results [37, 38] may relate specifically to object motion given that the stimuli used in these studies were smaller than the reported receptive field size of cells in the AOS. In contrast, our study likely engages the optic flow system, processed in the vermal lobules [41]. However, it should be pointed that the difference between the results of Cattaneo et al. [8] and the current experiment may also be due to the difference in stimulation technique (electromagnetic induction versus direct current). Specifically, tDCS can hyperpolarize and hypopolarize resting membrane potentials [48], whereas TMS induces action potentials.

Another consequence of the low spatial resolution of tDCS is that the current may spread beyond the structures of interest. In our case, the proximity of V1 may be problematic as one could argue that our results are may be driven by V1 activation. However, as shown in our model of the current distribution, the current spread to V1 as well as V5 is minimal (Fig. 2). Furthermore, from a psychophysical point of view, activity in V1 alone would not help the participants in our task because of the so-called aperture problem [49, 50, 51, 52]). That is, the receptive fields of V1 neurons are too small to process global motion in a large motion array such as this one. Therefore, to alter the processing of optic flow you would need to affect a part of the motion processing system with much larger receptive fields. The nearest structure that processes motion with receptive fields of that size is the cerebellum. Therefore, we do not think that our results are related to tDCS induced changes in V1 or MT.

A potential hypothesis for why we found an effect of anodal tDCS only with left-right motion discrimination is that with the limited penetration capability of tDCS, the current may have only affected a neuronal population that respond to left-right motion. Evidence for the segregation of neuronal populations responding to different motion directions are abundant in the visual system, including the accessory optic system that is known to process optic flow and project to the cerebellum [16]. For instance, in various species, optic flow neurons in the cerebellum are organized along a parasagittal plane, alternating different motion orientations [53, 54, 55]. Therefore, our results could be indicative of a similar direction selective segregation of cells within the cerebellum such that tDCS was only able to penetrate deep enough to affect neuronal populations involved in processing LR but not UD motion. Further investigations using molecular markers in non-human animals or electrophysiological recordings can help clarify this issue.

The polarity of tDCS applied to cortical structures is known to induce systematic changes in excitability. That is, anodal (+) current is thought to be excitatory whereas cathodal (−) is inhibitory. In the current study, we found both anodal and cathodal stimulation increased performance of left-right motion discrimination, although only the anodal current statistically improved performance relative to sham. These effects were apparent within the first ten minutes of stimulation. This non-specific effect of polarity on the cerebellum has been noted elsewhere [19, 56]. Specifically, in a recent meta analysis examining the effects of cerebellar tDCS, van Dun and colleagues [19] noted that although there was clear evidence that cerebellar tDCS could influence cognitive and motor processes, there were no consistent polarity specific effects found across studies. Primarily, the established notion that anodal and cathodal stimulation induces excitation and inhibition respectively cannot be applied to the cerebellum [57]. This is likely due to the physiological organization of the cerebellum. The cerebellum is more densely packed with neurons compared to the cortex and is highly convoluted. A corollary of this is that cells differ in orientation within a smaller area. Given that the effects of tDCS can be dependent upon the orientation of cells [58, 59], this likely explains the differences between cortical and cerebellar stimulation. It is, however, an open question that remains to be investigated.

Overall, the present study has several important findings. First, we showed that anodal stimulation of the midline cerebellum can improve optic flow perception during and post stimulation (‘offline’). This finding has clinical implications since it suggests that tDCS may be helpful with individuals that exhibit perceptual deficits relating to motion. Indeed, cerebellar tDCS has shown promising results with patients suffering from movement disorders [60]. Second, we also showed that, at least with tDCS, this effect may be limited to left-right motion. Based on the current study, future studies should investigate the duration of the improvement in motion perception as well as whether complex motion patterns (e.g., radial motion) are also influenced by cerebellar midline stimulation.

