## Supplemental Table 1 for "Cerebellar tDCS alters the perception of optic flow"

**Supplementary material**

Table 1. Change in accuracy relative to threshold for each participant in each condition in Experiment 1. The codes are as follows: UD = Up-down motion discrimination; LR = Left-right motion discrimination; S1 = 10 minutes stimulation; S2 = 20 minutes stimulation; Post = Post-stimulation; A = Anode, C = Cathode; S = Sham. As an example, UDS1A = accuracy for up-down motion after 10 minutes of stimulation of anodal stimulation.

| Participant ID | UDS1A | UDS2A | UDPostA | LRS1A | LRS2A | LRPostA |
| --- | --- | --- | --- | --- | --- | --- |
| 1 | -0.26 | 0.02 | 0.07 | 0.14 | 0.14 | 0.16 |
| 2 | 0.11 | 0.17 | 0.13 | 0.11 | 0.12 | 0.12 |
| 3 | -0.04 | 0.11 | 0.16 | 0.14 | 0.19 | 0.23 |
| 4 | 0.07 | 0.19 | 0.14 | 0.05 | 0.25 | 0.19 |
| 5 | 0.06 | 0.15 | 0.13 | 0.21 | 0.26 | 0.23 |
| 6 | -0.02 | -0.07 | 0.13 | 0.18 | 0.13 | 0.15 |
| 7 | 0.08 | 0.16 | 0.09 | 0.16 | 0.24 | 0.25 |
| 8 | 0.19 | 0.26 | 0.26 | 0.23 | 0.22 | 0.24 |
| 9 | -0.05 | 0.08 | 0.09 | 0.17 | 0.21 | 0.21 |
| 10 | 0.12 | 0.22 | 0.26 | 0.04 | 0.07 | 0.05 |
| 11 | 0.06 | 0.06 | 0.12 | 0.06 | -0.02 | 0.09 |
| 12 | 0.22 | 0.15 | 0.25 | 0.22 | 0.25 | 0.26 |
| 13 | 0 | 0.13 | 0.16 | -0.07 | 0.1 | 0.08 |
| 14 | 0.01 | 0.05 | 0.05 | 0.16 | 0.14 | -0.01 |
| 15 | -0.08 | 0.04 | 0.08 | 0.01 | -0.06 | 0.11 |
| 16 | 0.11 | 0.07 | 0.09 | 0 | -0.17 | 0.05 |

Table 1. Cont’d

| Participant ID | UDS1C | UDS2C | UDPostC | LRS1C | LRS2C | LRPostC |
| --- | --- | --- | --- | --- | --- | --- |
| 1 | 0.21 | 0.26 | 0.24 | 0.22 | 0.24 | 0.25 |
| 2 | 0.1 | 0.07 | 0.26 | -0.02 | 0.02 | 0.18 |
| 3 | -0.12 | -0.01 | 0.13 | -0.08 | -0.01 | 0 |
| 4 | 0.06 | 0.12 | 0.11 | 0.08 | 0.19 | 0.18 |
| 5 | 0.01 | 0.19 | -0.05 | 0.05 | 0.1 | 0.12 |
| 6 | 0.04 | 0.16 | 0.08 | -0.08 | -0.07 | -0.12 |
| 7 | 0.11 | 0.11 | 0.2 | -0.03 | 0.07 | 0.02 |
| 8 | 0.05 | 0.15 | 0.21 | 0.02 | 0.08 | 0.11 |
| 9 | 0.13 | 0.18 | 0.2 | 0.03 | 0.1 | 0.14 |
| 10 | 0.08 | 0.15 | 0.21 | 0.09 | 0.12 | 0.03 |
| 11 | 0.09 | 0.07 | 0.16 | 0.05 | 0.04 | -0.03 |
| 12 | 0.04 | 0.15 | 0.01 | -0.1 | -0.08 | 0.05 |
| 13 | 0.14 | 0.1 | 0.12 | -0.05 | -0.01 | -0.03 |
| 14 | -0.11 | -0.08 | 0.01 | 0.1 | 0.14 | 0.15 |
| 15 | 0.09 | 0.18 | 0.23 | 0.15 | 0.28 | 0.21 |
| 16 | -0.09 | 0.04 | 0.04 | 0.14 | 0.14 | 0.14 |

Table 1. Cont’d

| Participant ID | UDS1S | UDS2S | UDPostS | LRS1S | LRS2S | LRPostS |
| --- | --- | --- | --- | --- | --- | --- |
| 1 | 0.07 | 0.18 | 0.17 | 0 | 0.09 | 0.02 |
| 2 | 0 | 0.07 | -0.14 | 0 | 0.08 | -0.15 |
| 3 | 0.02 | 0.14 | 0.04 | 0.04 | 0.02 | -0.21 |
| 4 | 0.15 | 0.16 | 0.21 | 0.24 | 0.28 | 0.28 |
| 5 | -0.01 | 0.09 | -0.1 | -0.21 | -0.02 | -0.11 |
| 6 | -0.07 | 0.22 | 0.14 | -0.12 | 0.04 | -0.16 |
| 7 | 0.13 | 0.19 | 0.2 | -0.05 | 0.03 | 0.08 |
| 8 | 0.22 | 0.09 | 0.28 | -0.07 | 0.02 | 0.11 |
| 9 | -0.01 | 0.06 | -0.04 | -0.11 | -0.01 | -0.04 |
| 10 | 0.08 | 0.13 | 0.24 | 0.03 | 0.18 | 0.25 |
| 11 | 0.03 | 0.05 | -0.02 | -0.03 | 0.16 | 0.18 |
| 12 | 0.06 | 0.03 | 0.03 | 0.12 | -0.02 | -0.01 |
| 13 | -0.02 | 0.17 | 0.02 | -0.24 | -0.15 | -0.17 |
| 14 | -0.02 | 0.02 | 0.03 | 0.06 | 0 | 0.06 |
| 15 | -0.14 | -0.04 | 0 | 0.1 | -0.02 | 0.08 |
| 16 | 0.09 | 0.13 | 0.16 | -0.06 | 0.07 | 0.11 |
